# Activation of Macrophages by Extracellular Vesicles Derived from *Babesia*-infected Red Blood Cells

**DOI:** 10.1101/2024.02.19.581030

**Authors:** Biniam Hagos, Ioana Brasov, Heather Branscome, Fatah Kashanchi, Choukri Ben Mamoun, Robert E. Molestina

## Abstract

*Babesia microti* is the primary cause of human babesiosis in North America. Despite an emergence of the disease in recent years, the pathogenesis and immune response to *B. microti* infection remain poorly understood. Studies in laboratory mice have shown a critical role for macrophages in the elimination of parasites and infected red blood cells (iRBCs). Importantly, the underlying mechanisms that activate macrophages are still unknown. Recent evidence identified the release of extracellular vesicles (EVs) from *Babesia* iRBCs. EVs are spherical particles released from cell membranes under natural or pathological conditions that have been suggested to play roles in host-pathogen interactions among diseases caused by protozoan parasites. The present study examined whether EVs released from cultured *Babesia* iRBCs activated macrophages resulting in changes in the secretion of cytokines. An analysis of vesicle size in EV fractions from *Babesia* iRBCs showed diverse populations in the <100 nm size range compared to EVs from uninfected RBCs. Uptake of EVs released from *B. microti*-iRBCs was observed in macrophages *in vitro*. In addition, incubation of macrophages with EVs isolated from *Babesia* iRBC culture supernatants resulted in the activation of NF-κB and modulation of pro-inflammatory cytokines. These results support a role for EVs in the activation of macrophage functions and provide new insights into the mechanisms involved in the induction of the innate immune response during babesiosis.

## INTRODUCTION

Babesiosis is an emerging tickborne disease in the United States caused by intraerythrocytic protozoan parasites of the genus *Babesia* (Gray et al., 2019). *Babesia microti* is the primary causative agent for most human cases of babesiosis. Transmission occurs primarily by Ixodes tick vectors, with less common routes of human-to-human transmission including pregnancy and blood transfusion (Krause, 2019; Vannier et al., 2015). The latter is of concern given the fact that babesiosis is the most frequently reported transfusion-transmitted parasitic infection in the U.S. (Gubernot et al., 2009; Leiby, 2011; Lobo et al., 2013). Human babesiosis is usually asymptomatic in immunocompetent populations, or results in mild symptoms that resolve within a few days. Immunocompromised individuals may experience severe symptoms of acute anemia, thrombocytopenia, organ failure, and even death (Krause, 2019; Vannier et al., 2015).

Despite the emergence of human cases of babesiosis in recent years, the pathogenesis and immune response to *B. microti* infection remain poorly understood. Splenectomized or innately asplenic mice are highly susceptible to infection, whereas infected immunocompetent mice and hamsters display significantly enlarged spleens due to increased numbers of macrophages (Coleman et al., 2005; Djokic et al., 2018a). Macrophage depletion results in elevated parasitemia and mortality in mice, highlighting the contribution of macrophages in the elimination of parasites and parasitized RBCs during babesiosis (Terkawi et al., 2015). Importantly, the effector parasite molecules that trigger the innate immune response to *B. microti* are still unknown.

Contrary to *Plasmodium* which develops inside a parasitophorous vacuole (PV) following RBC invasion, *B. microti* forms a transient PV upon invasion which eventually disintegrates as the parasite develops in the RBC cytoplasm (Rudzinska, 1976; Rudzinska et al., 1976). Giemsa-stained blood smears of *B. microti*-infected RBCs show different morphological changes such as ring-shaped forms, membranous extensions protruding from rings, and the less common tetrad forms (Thekkiniath et al., 2019). The membranous extensions have been described as dendrite-like tubulovesicular structures or tubes of vesicles (TOVs) (Thekkiniath et al., 2019). These TOVs originate by the vesiculation of the *B. microti* plasma membrane followed by an interlacement of connected vesicles within the RBC cytoplasm. It has been postulated that this system of vesicles extending from the parasite into the RBC represents a novel mechanism of protein trafficking and export (Thekkiniath et al., 2019). Importantly, the vesicle-mediated protein export is not only restricted to the host RBC but also to the extracellular environment, as shown by the detection of *B. microti* immunodominant antigens in EVs isolated from the plasma of infected mice (Beri et al., 2022; Thekkiniath et al., 2019). A recent report by Beri et al. identified EVs released from RBCs infected *in vitro* with *B. divergens* (Beri et al., 2022).

We posit that the recently described *B. microti* protein export system is a mechanism by which parasite antigens enclosed within EVs released to the extracellular environment participate in cell-to-cell communication, similar to mechanisms reported with malaria (Couper et al., 2010; Mantel et al., 2016; Mantel et al., 2013; Mbagwu et al., 2019). When macrophages function as recipients of EVs from *B. microti* iRBCs, changes in the modulation of cytokines with key roles in the host innate immune response to the parasite occur. To address this hypothesis, this study examined cytokine responses in macrophages exposed to EVs derived from *B. microti* iRBCs. We also assessed the diversity of vesicle populations found in iRBC-derived EV fractions by size distribution analysis and examined EV uptake by macrophages. Results from this study provide insights into the mechanisms of intercellular communication between *Babesia* and macrophages which are plausibly critical to the induction of the innate immune response in the mammalian host.

## MATERIALS AND METHODS

### Babesia microti isolate

*Babesia microti* GI (ATCC^®^ PRA-398^TM^) was originally isolated from blood obtained from a human case of babesiosis in Nantucket, Massachusetts, USA, in 1983 (Gray et al., 2002; Piesman et al., 1986). The isolate was maintained by in vivo propagation in Syrian hamsters (Stock HsdHan:AURA, Inotiv, Indianapolis, IN) according to published protocols (Cullen and Levine, 1987; Oz and Hughes, 1996) and procedures approved by the ATCC^®^ Institutional Animal Care and Use Committee.

### Short-term *in vitro* culture of *B. microti*

Blood was collected from a *B. microti*-infected hamster at 25% parasitemia and leukocytes were depleted by passing the sample through an Acrodisc^®^ white blood cell syringe filter (Pall Biotech, Westborough, MA). Infected blood was diluted to 5% parasitemia with uninfected human blood (Interstate Blood Bank, Philadelphia, PA) at 50% hematocrit. Erythrocyte cultures were established in 6 well plates using HL-1 medium (Lonza, Basel, Switzerland) supplemented with 20% human serum type A^+^ (Interstate Blood Bank), 1% (v/v) HB 101 (Irvine Scientific, Santa Ana, CA), 2 mM L-glutamine (ATCC^®^, Manassas, VA), 2X hypoxanthine/thymidine solution, 1X antibiotic/antimycotic solution, and 100 μg/ml gentamicin solution (ThermoFisher, Waltham, MA). Cultures were incubated at 37°C under 2% O_2_/5% CO_2_/93% N_2_ atmospheric conditions for up to 72h (Abraham et al., 2018). Parasitemia was checked daily by microscopic examination of Giemsa-stained blood smears.

### Macrophage cultures

The THP-1 human monocytic cell line ATCC^®^ TIB-202^TM^ and the NF-κB reporter human monocytic cell line ATCC^®^ TIB-202-NFkB-LUC2^TM^ were maintained at 37°C with 5% CO_2_ in RPMI-1640 medium supplemented with 10% FBS, 100 U/ml penicillin, and 100 µg/ml streptomycin. Monocyte-derived macrophages were derived by treating the cell lines with 100 ng/ml of phorbol 12-myristate 13-acetate (PMA; Sigma, St. Louis, MO) for 48 h followed by a 24 h incubation in medium alone before experiments. All medium components were obtained from ATCC^®^ and cell lines were tested for the absence of bacterial contamination using BacT/ALERT (BioMérieux, Durham, NC) and mycoplasma contamination by Hoechst DNA stain, culture, and PCR.

### Isolation of EVs from RBC culture supernatants

uRBC and *B. microti* iRBC cultures were established in 6-well plates using the conditions described above. Supernatants (15 ml) were collected after 24 and 48 h of incubation and centrifuged at 500 *g* for 10 min to pellet RBCs. Samples were subsequently subjected to stepwise centrifugations at 2,000 g for 10 min (2K), 10,000 g for 40 min (10K), 100,000 g for 90 min (100K), and 167,000 g for 16 h (167K) (Barclay et al., 2019; DeMarino et al., 2018). EV pellets were resuspended in 200 μl of PBS and stored at −80°C before use. In separate experiments, EVs were enriched from RBC culture supernatants using ExoMax^TM^ according to manufacturer’s instructions (System Biosciences, Palo Alto, CA). EVs were examined for protein concentration using BCA and by Western blots as described below.

### Western blots

Proteins in EV samples were resolved by SDS-PAGE and transferred to polyvinylidene difluoride (PVDF) membranes. PVDF membranes were blocked in PBS with 3% non-fat milk powder and 0.05% Tween 20 for 1 h. Membranes were then probed with 1:500 dilutions of rabbit polyclonal antibodies raised against host parasite antigen BmIPA48 (Magni et al., 2020). Primary antibody binding was detected by a goat-anti-rabbit antibody conjugated to horseradish peroxidase (HRP) (1:2000 dilution; ThermoFisher). Signals were detected using a chemiluminescence substrate and the Azure c600 Imaging System (Azure Biosystems, Dublin, CA).

### Quantification of EVs and analysis of EV size during infection

The concentration (particles/ml) and diameter of EVs isolated from culture supernatants were examined by Nanoparticle Tracking Analysis (NTA) using the Nanosight NS300 instrument (Malvern Panalytical, Westborough, MA). Each sample was analyzed in triplicate with equipment settings remaining constant between readings. Comparisons between EV fractions obtained from RBC culture supernatants during the 24 to 48 h time frame were performed to evaluate possible changes in the numbers and size distributions of EVs released during infection. Analysis of samples from uninfected RBCs served as controls to determine potential increases in EV secretion as a result of *B. microti* infection.

### NF-κB Activation Assay

Monocyte-derived macrophages of the ATCC^®^ TIB-202-NFkB-LUC2™ cell line were cultured in 12-well plates at 2 x 10^6^ cells/well and incubated for 24 h at 37°C/5% CO_2_ with increasing concentrations of EVs from uRBC or *B. microti* iRBC culture supernatants. Macrophages treated with 5 μg/ml of LPS for 24 h were used as positive controls. Following incubations, culture supernatants were frozen at −s80°C and macrophages were processed according to the Promega E1500 Luciferase Assay System protocol (Promega, Madison, WI). Cells were lysed and incubated with D-Luciferin substrate for 5 min in white Lumitrac™ plates (Greiner Bio-one, Monroe, NC). Luminescence was analyzed using a SpectraMax M5 system connected to SoftMax Pro version 8.0 software (Molecular Devices, San Jose, CA).

### Cytokine arrays

Macrophage supernatants were thawed on ice and assayed for the presence of 40 cytokines using Proteome Profiler Antibody Arrays (R&D Systems, Minneapolis, MN). Procedures were followed according to manufacturer’s instructions. Chemiluminescent signals from the arrays corresponding to the different cytokines were detected using the Azure^TM^ c600 Imaging System (Azure Biosystems). Densitometric analyses of cytokine spots were performed using the ImageJ software (https://ij.imjoy.io/). Integrated densitometric values (IDV) were normalized to spots corresponding to PBS negative controls on the arrays and the numerical data was plotted using GraphPad Prism software version 8.0.

### Uptake of EVs by macrophages

EVs isolated in 167K fractions from RBC supernatants were labeled with BODIPY 493/503 (ThermoFisher). The dye was prepared in DMSO at a concentration of 1 mM. Five µL of the dye stock solution were mixed with approximately 10^7^ EVs in 100 µL of PBS. The EV suspension was incubated at 37°C for 30 min in the dark and loaded into spin columns packed with G-10 Sephadex. Excess unincorporated dye was removed by centrifugation of the spin columns at 500 *g* for 2 minutes. The efficiency of EV labeling and quantification of EVs was performed by NTA as described above. Monocyte-derived macrophages of the ATCC^®^ TIB-202^TM^ cell line cultured in 8-well chamber slides at 2 x 10^4^ cells/well were incubated at 37°C/5% CO_2_ for 0.5, 1, 2, and 3 h with approximately 2 x 10^6^ labeled EVs/well. Cells were fixed in 4% paraformaldehyde for 10 min, washed in PBS, and visualized under 640X magnification using a Zeiss Axioscope fluorescence microscope connected to a digital camera. Digital microscopic images were captured using Zen Imaging Software (Zeiss, Oberkochen, Germany).

### Data analysis

Data from NF-κB luciferase assays were collected from three experiments. In each experiment, luminescence measurements were performed in duplicate for each sample (Fig. 4). Data was subsequently analyzed using GraphPad Prism 8 (GraphPad Software, Inc., San Diego, CA, USA) to calculate means and standard error of the means (SEM). Where indicated, results were subjected to analysis of variance followed by the Tukey’s multiple-comparison test. A *P* value of <0.05 was used to determine statistical significance.

## RESULTS

### In vitro culture of B. microti

A continuous *in vitro* model system for *B. microti* is currently unavailable, however, small scale studies are possible using short term cultures. In our studies, blood was collected from *B. microti*-infected hamsters at 25% parasitemia and diluted to 5% parasitemia with uninfected human blood at 50% hematocrit. RBC cultures were established in 6 well plates and incubated at 37°C under 2% O_2_/5% CO_2_/93% N_2_ atmospheric conditions as described (Abraham et al., 2018). As shown in Fig. 1A, an increase in parasitemia to approximately 12% and 14% was observed after 24h and 48 h of inoculation, respectively, followed by a decrease after 72 h. Microscopic analysis of in vitro cultures showed singly and multiply infected cells with merozoites and ring stages of the parasite (Fig. 1B, arrows).

**Fig. 1.**
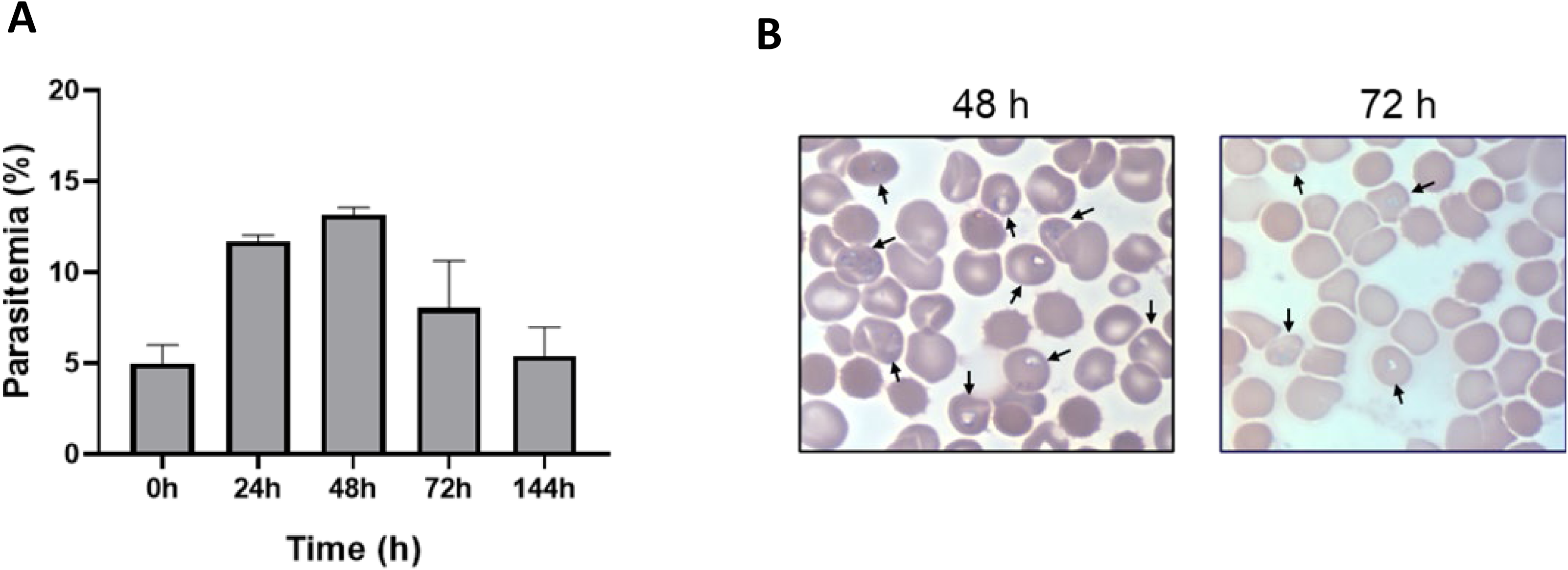
Short-term *in vitro* culture of *B. microti*. A, parasitemia of iRBC cultures was determined by microscopy. Columns represent means+SEM of three replicates from a representative experiment. B, representative image of intracellular stages of *B. microti* in iRBCs (arrows). Bar, 6 μm).

### Analysis of EV fractions isolated from iRBC supernatants

To examine the release of EVs from *B. microti* iRBCs cultured in vitro, supernatants were collected at different times of infection and subjected to stepwise centrifugations (Fig. 2A). Western blots detected the parasite antigen BmIPA48 in EV pellets collected following centrifugations of 2,000 g (2K), 10,000 g (10K), 100,000 g (100K), and 167,000 g (167K) (Fig. 2B). The majority of BmIPA48 was found associated with EVs from 2K and 10K pellets which represent large vesicles (>1 μm) and microvesicles (0.1-1 μm), respectively. The protein was detected to a significantly lesser degree in EVs from 100K pellets but markedly present in EVs from 167K fractions which are expected to harbor exosome sized vesicles (30-150 nm) (Fig. 2B).

**Fig. 2.**
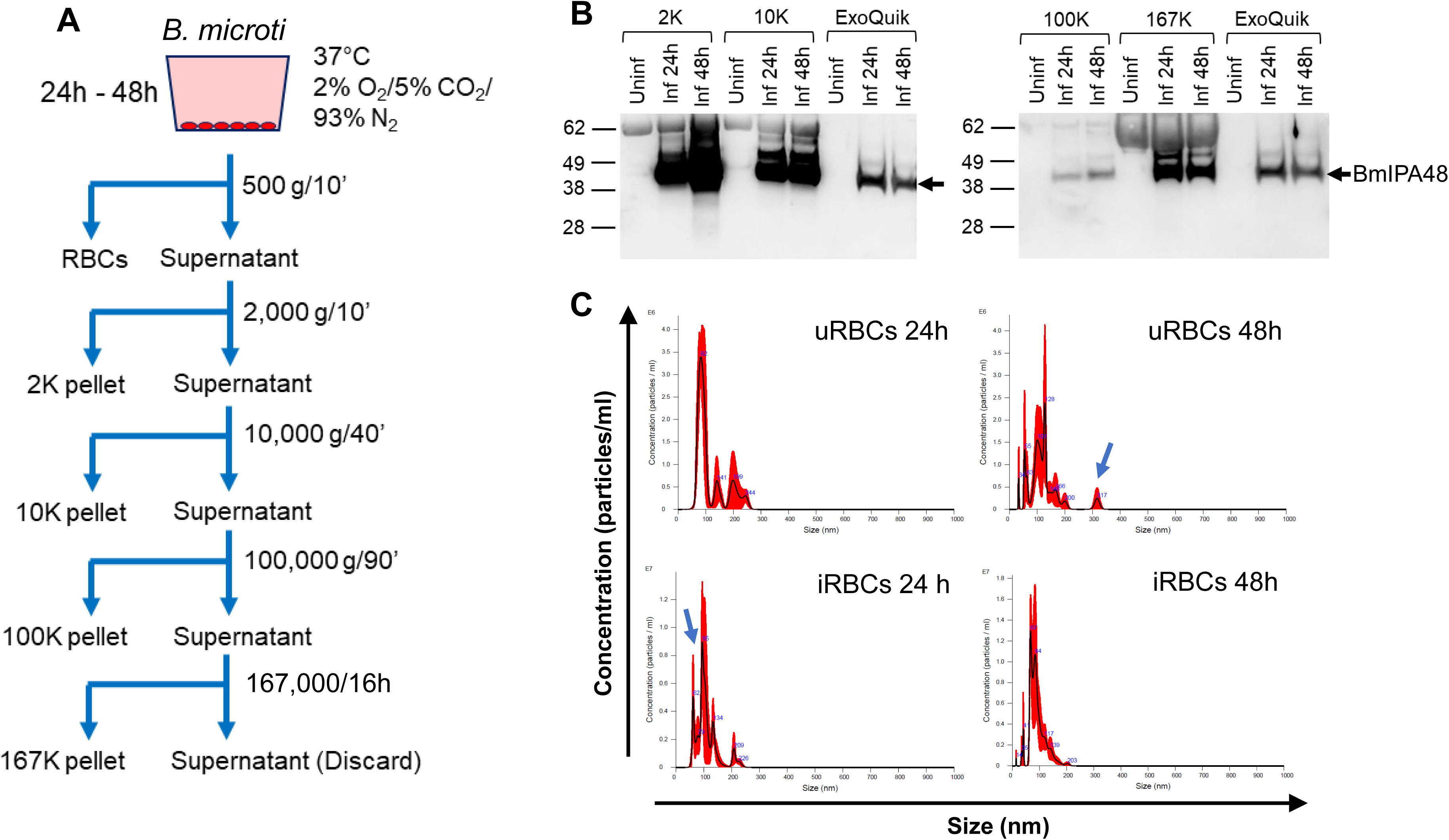
Analysis of EV fractions isolated from RBC culture supernatants. A, schematic for the isolation of EVs by sequential centrifugation. B, BmIPA48 Western blots in EVs isolated from uRBC (Uninf) and iRBC (Inf) supernatants. C, size distribution of EVs isolated in 167K pellets by NTA. Arrows depict distinct EV populations observed in uninfected and infected samples.

Previous immunoelectron microscopy analyses of *B. microti*-infected RBCs and plasma from infected mice identified parasite proteins associated with vesicles of ∼100 nm in diameter (Thekkiniath et al., 2019). Thus, we examined the size distribution of EVs found in our 167K fractions by NTA to determine if vesicles of similar diameter were detected. As shown in Fig. 2C, the sizes of vesicles in uRBC and *Babesia* iRBC EVs isolated from 24 h culture supernatants ranged from 60 to 250 nm. The mean diameter of uRBC EVs was 114.4±15.9 nm, with peaks observed at 82, 141, 199, and 244 nm. *Babesia* iRBC EVs were similar in diameter to uRBC EVs measuring 110.8±1.8 nm, however, the size range included an additional smaller peak at 62 nm and peaks detected at 79, 95, 134, 209, and 226 nm (Fig. 2C). Interestingly, the mean concentration of EVs was twofold higher in 167K fractions of 24 h iRBC cultures compared to uRBC cultures (3.45 x 10^8^ versus 1.53 x 10^8^ particles/ml, respectively). This difference increased to fourfold after 48 h of culture (1.16 x 10^8^ particles/ml in uRBC versus 5.03 x 10^8^ particles/ml in *Babesia* iRBC cultures). In addition, a widening in vesicle size ranges from 34 to 320 nm and 14 to 203 nm were noticeable in 48 h cultures of uRBCs and iRBCs, respectively. The highest concentrations of EVs in these 167K fractions were found in the 55-130 nm range for uRBCs and 40-85 nm range for iRBCs (Fig. 2C).

### Macrophage uptake of EVs isolated from RBC culture supernatants and hamster plasma

To investigate whether macrophages were able to internalize EVs from *Babesia* iRBCs, monocyte-derived macrophages of the ATCC^®^ TIB-202^TM^ cell line were exposed to BODIPY-labeled vesicles present in the 167K fractions. As shown in Fig. 3, clusters of intracellular fluorescence and fluorescent punctate signals were observed in macrophages after 90 min of exposure to BODIPY labeled EVs from uRBC and iRBC cultures. Fluorescent clusters were generally detected inside macrophages indicative of uptake internalization of BODIPY-labeled EVs (Fig. 3A-B, arrows), while punctate signals were primarily localized at the macrophage cell membranes (Fig. 3A-B, arrowheads). Exposure of cells to dye-only control solution did not result in intracellular fluorescent staining (Data not shown).

**Fig. 3.**
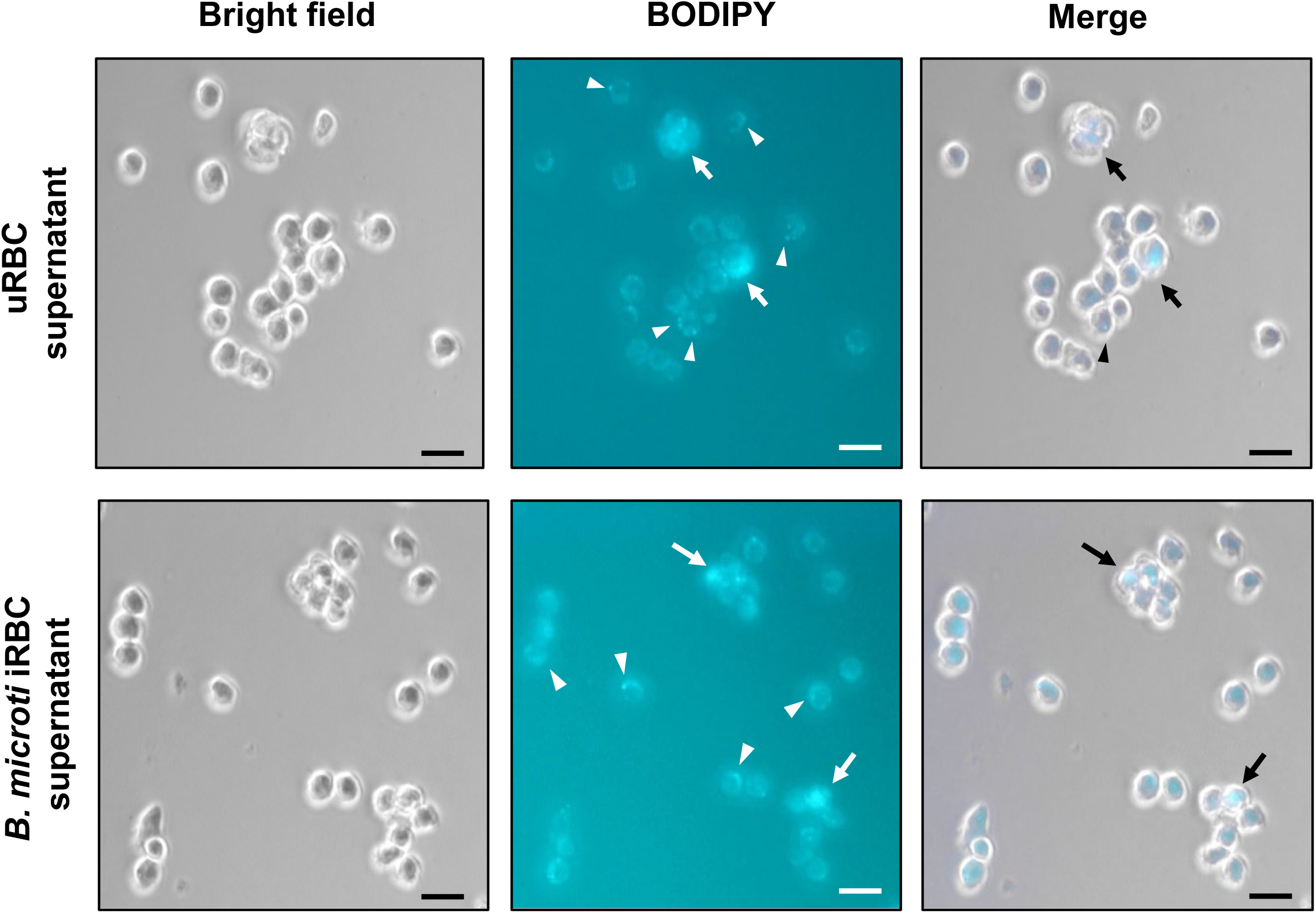
Macrophage uptake of EVs isolated from uRBC supernatants (A) and *B. microti* iRBC supernatants (B). EVs present in 167K fractions were labeled with BODIPY dye and incubated with monocyte-derived macrophages (ATCC^®^ TIB-202^TM^) for 90 min. Arrows show internalization of BODIPY-labeled EVs. Arrowheads show localization of BODIPY-labeled EVs at the macrophage cell membranes. Bar, 15 μm.

### Activation of NF-κB by EVs isolated from iRBC supernatants

The effect of EV treatment on the transcription factor NF-κB was examined in macrophages by luciferase reporter assays. Monocyte-derived macrophages (ATCC^®^ TIB-202-NFκB-LUC2™) were incubated for 24 h with different protein concentrations of EVs found in 167K fractions. As shown in Fig. 4A, a significant increase in NF-κB-dependent luciferase activity was detected in response to 50 μg/ml of EVs from *Babesia* iRBCs compared to uRBC controls. A modest increase was observed at the lower EV iRBC concentration of 5 μg/ml. In parallel to the increase in NF-κB activity, increases in pro-inflammatory mediators were detected in the supernatants of macrophages treated with 50 μg/ml of iRBC EVs as compared to uRBC EVs (Fig. 4B). Among specific cytokines known to be regulated by NF-κB (Liu et al., 2017) we found stimulation of GM-CSF, IFN-γ, IL-1β, IL-6, IL-8, IL-12, and IL-18 in response to treatment with iRBC EVs.

**Fig. 4.**
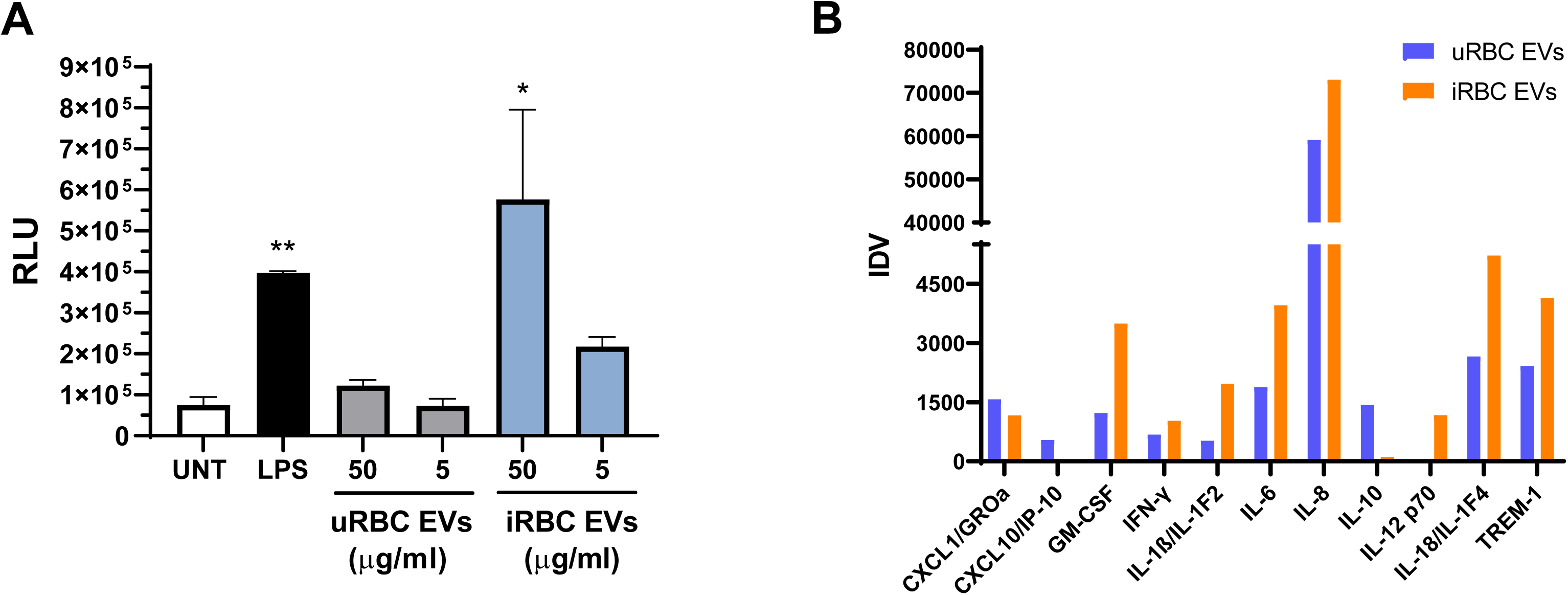
Macrophage NF-κB activity and cytokine production in response to EVs from *Babesia* iRBCs. A, luciferase activity was measured in macrophages treated for 24 h with different protein concentrations of 167K EV fractions isolated from uRBC or iRBC supernatants. UNT, untreated; LPS, macrophages treated with 5 μg/ml of LPS. Bars represent means+SEM of three experiments. **, *P*<0.01 compared to UNT; *, *P*<0.05 compared to 50 μg/ml of uRBC EVs. B, production of cytokines in macrophages treated with EVs for 24 h. Culture supernatants were examined by the Proteome Profiler Antibody Array kit (R&D Systems). Densitometric analysis of protein spots from a representative experiment show changes in 11 cytokines in response to 50 μg/ml of iRBC EVs (orange bars) compared to uRBC EVs (blue bars).

## DISCUSSION

There has been a significant increase in EV research in the past decade, particularly on the role of EVs in host-pathogen interactions among diseases caused by protozoan parasites (de Souza and Barrias, 2020; Gavinho et al., 2018). Silverman et al. reported the release of exosomes by *Leishmania* promastigotes in response to changes in temperature and pH (Silverman et al., 2010a). Uptake of *Leishmania* exosomes by macrophages stimulated IL-8 production (Silverman et al., 2010b). In Chagas disease, the release of EVs occurs in epimastigote and trypomastigote stages of *Trypanosoma cruzi*, causing the induction of proinflammatory cytokines and nitric oxide by macrophages (Nogueira et al., 2015). In *Plasmodium*, EVs are elevated in the plasma of malaria patients in proportion to disease severity (Nantakomol et al., 2011). When purified from the plasma of malaria-infected mice, EVs induced potent activation of macrophages via Toll-like receptor (TLR) signaling (Couper et al., 2010). EVs released from *in vitro* cultures of *Plasmodium*- infected RBCs were shown to activate macrophages to produce cytokines and stimulate chemotaxis in neutrophils (Mantel et al., 2013). In addition to playing a role in the pathology of malaria, *P. falciparum* takes advantage of RBC microvesicle pathways to induce the release of EVs from infected cells to mediate intercellular communication between RBCs, facilitate horizontal transfer of nucleic acids, and regulate parasite density and production of high numbers of gametocytes *in vitro* (Regev-Rudzki et al., 2013).

Prior work from Thekkiniath et al. (Thekkiniath et al., 2019) provided us with a blueprint to study the basis of EV release from parasitized RBCs. The authors suggested that *B. microti*- derived vesicles are actively exported during parasite development with little, if any, involvement from host microvesicle pathways. Their findings were deduced from studies in EV pellets obtained by ultracentrifugation from murine plasma (Thekkiniath et al., 2019). Using *Babesia in vivo* models to study biological activities of parasite-derived EVs carries certain limitations given that EVs in plasma can originate from erythrocytes, leukocytes, platelets, and endothelial cells (Brahmer et al., 2019; Yanez-Mo et al., 2015). To minimize the effects of EVs from other cellular sources, the present study approached the isolation of EVs from *in vitro* cultured RBCs. Of note, a continuous *in vitro* model system for *B. microti* is currently unavailable, however, small scale studies are possible using short term cultures.

*B. microti* iRBCs cultured *in vitro* release EVs harboring the parasite antigen BmIPA48 as shown by ultracentrifugation of vesicle pellets from culture supernatants and immunoblot detection. The presence of BmIPA48 in vesicles was also detected by enrichment of EVs from *B. microti* iRBC culture supernatants with the polymer-based reagent ExoQuick. Whether BmIPA48 is exported in close association with EV membranes or inside EVs remains to be determined. The latter scenario is more likely given the absence of a GPI-anchor motif or a transmembrane domain in BmIPA48 (Silva et al., 2016). A comparison of EV size distributions found in 167K fractions of 24 h uRBC and *Babesia* iRBC cultures showed notable differences, particularly among EVs <100 nm in diameter. The detection of specific parasite-derived vesicles among these distinct EV populations will require further fractionation experiments using gradient centrifugation or size-exclusion chromatography. These approaches combined with proteomics will help us gain a better understanding of the protein composition and biogenesis of vesicles secreted from *Babesia* iRBCs to the extracellular environment. The finding that the overall concentrations of EVs in 167K pellets were higher as a result of infection has also been observed with *B. divergens* (Beri et al., 2022), suggesting that *Babesia* infection modulates microvesicle generation in RBCs as reported with *Plasmodium* (Mantel et al., 2013).

Macrophage uptake of EVs was indistinguishable between vesicles isolated from uRBC and iRBC cultures. However, EVs isolated from iRBCs induced a significant increase in macrophage NF-κB activity and concomitant production of cytokines compared to EVs from uRBCs. Increases in pro-inflammatory cytokines are a hallmark of acute babesiosis in mouse studies (Djokic et al., 2018a; Djokic et al., 2018b; Skariah et al., 2017; Terkawi et al., 2015). The observed NF-κB and cytokine responses to iRBC EVs in macrophages warrant future studies to determine to what extent *Babesia*-derived EVs participate in the innate immune response to infection (Djokic et al., 2018a).

In summary, the present study provides evidence supporting that *B. microti* infection induces the release of EVs from RBCs and that EVs released from parasite iRBCs cause phenotypic changes in macrophages. The identification of the TOV system as a protein export mechanism is a noteworthy discovery in *Babesia* biology; however, many questions remain that need to be addressed (Fig. 5): 1) does *B. microti* manipulate RBC microvesicle pathways to induce the release of EVs to the extracellular environment? 2) is the extent of EV release dependent on parasite growth and egress? 3) to what degree do parasite-derived EVs participate in cell-to-cell communication resulting in phenotypic changes of resting recipient cells? 4) what are the parasite-derived protein cargoes encapsulated within EVs? 5) are parasite antigens transferred from EVs to recipient cells, and if so, does this transfer result in changes in signal transduction pathways? 6) do parasite antigens enclosed within EVs engage macrophage Toll-like receptors (TLRs) and drive the cytokine response? 7) how critical is the EV-dependent modulation of macrophage functions in the innate immune response to the parasite? Elucidating the roles of host and parasite factors involved in vesicle-mediated cell-to-cell communication and the modulation of host biological activities by *Babesia*-derived EVs will contribute to a better understanding of the mechanisms governing intracellular parasitism in babesiosis.

**Fig. 5.**
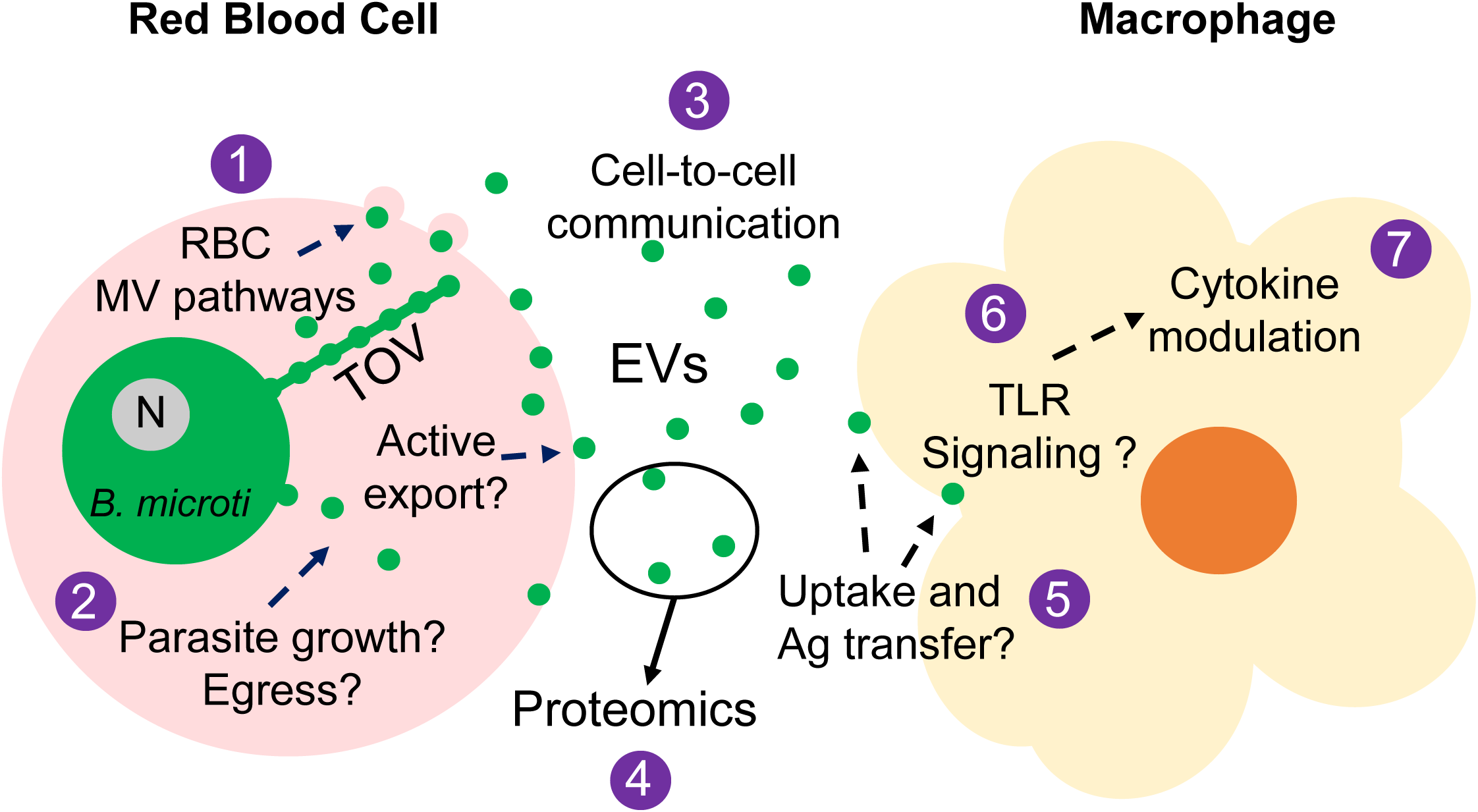
Plausible biological role of *B. microti* iRBC EVs in babesiosis. See text for details. Bm, B. microti; N, parasite nucleus; MV, microvesicle; TOV, tubes of vesicles (Thekkiniath et al., 2019).

## ACKNOWLEDGMENTS

This work was supported by the ATCC^®^ Internal Research and Development Program.

## REFERENCES

Abraham, A., Brasov, I., Thekkiniath, J., Kilian, N., Lawres, L., Gao, R., DeBus, K., He, L., Yu, X., Zhu, G., Graham, M.M., Liu, X., Molestina, R., Ben Mamoun, C., 2018. Establishment of a continuous in vitro culture of *Babesia duncani* in human erythrocytes reveals unusually high tolerance to recommended therapies. J Biol Chem 293, 19974–19981.

Barclay, R.A., Khatkar, P., Mensah, G., DeMarino, C., Chu, J.S.C., Lepene, B., Zhou, W., Gillevet, P., Torkzaban, B., Khalili, K., Liotta, L., Kashanchi, F., 2019. An Omics Approach to Extracellular Vesicles from HIV-1 Infected Cells. Cells 8.

Beri, D., Rodriguez, M., Singh, M., Liu, Y., Rasquinha, G., An, X., Yazdanbakhsh, K., Lobo, C.A., 2022. Identification and characterization of extracellular vesicles from red cells infected with *Babesia divergens* and *Babesia microti*. Front Cell Infect Microbiol 12, 962944.

Brahmer, A., Neuberger, E., Esch-Heisser, L., Haller, N., Jorgensen, M.M., Baek, R., Mobius, W., Simon, P., Kramer-Albers, E.M., 2019. Platelets, endothelial cells and leukocytes contribute to the exercise-triggered release of extracellular vesicles into the circulation. J Extracell Vesicles 8, 1615820.

Coleman, J.L., LeVine, D., Thill, C., Kuhlow, C., Benach, J.L., 2005. *Babesia microti* and *Borrelia burgdorferi* follow independent courses of infection in mice. The Journal of infectious diseases 192, 1634–1641.

Couper, K.N., Barnes, T., Hafalla, J.C., Combes, V., Ryffel, B., Secher, T., Grau, G.E., Riley, E.M., de Souza, J.B., 2010. Parasite-derived plasma microparticles contribute significantly to malaria infection-induced inflammation through potent macrophage stimulation. PLoS pathogens 6, e1000744.

Cullen, J.M., Levine, J.F., 1987. Pathology of experimental *Babesia microti* infection in the Syrian hamster. Laboratory animal science 37, 640–643.

de Souza, W., Barrias, E.S., 2020. Membrane-bound extracellular vesicles secreted by parasitic protozoa: cellular structures involved in the communication between cells. Parasitol Res 119, 2005–2023.

DeMarino, C., Pleet, M.L., Cowen, M., Barclay, R.A., Akpamagbo, Y., Erickson, J., Ndembi, N., Charurat, M., Jumare, J., Bwala, S., Alabi, P., Hogan, M., Gupta, A., Noren Hooten, N., Evans, M.K., Lepene, B., Zhou, W., Caputi, M., Romerio, F., Royal, W., 3rd, El-Hage, N., Liotta, L.A., Kashanchi, F., 2018. Antiretroviral Drugs Alter the Content of Extracellular Vesicles from HIV-1-Infected Cells. Scientific reports 8, 7653.

Djokic, V., Akoolo, L., Parveen, N., 2018a. *Babesia microti* Infection Changes Host Spleen Architecture and Is Cleared by a Th1 Immune Response. Front Microbiol 9, 85.

Djokic, V., Primus, S., Akoolo, L., Chakraborti, M., Parveen, N., 2018b. Age-Related Differential Stimulation of Immune Response by *Babesia microti* and *Borrelia burgdorferi* During Acute Phase of Infection Affects Disease Severity. Front Immunol 9, 2891.

Gavinho, B., Rossi, I.V., Evans-Osses, I., Inal, J., Ramirez, M.I., 2018. A new landscape of host-protozoa interactions involving the extracellular vesicles world. Parasitology 145, 1521–1530.

Gray, E.B., Niknafs, A.M., Herwaldt, B.L., 2019. Surveillance for Babesiosis - United States, 2017. Annual Summary. https://www.cdc.gov/parasites/babesiosis/resources/babesiosis_surveillance_summary_2017b.pdf. Accessed 5 January 2020.

Gray, J., von Stedingk, L.V., Gurtelschmid, M., Granstrom, M., 2002. Transmission studies of *Babesia microti* in Ixodes ricinus ticks and gerbils. Journal of clinical microbiology 40, 1259–1263.

Gubernot, D.M., Nakhasi, H.L., Mied, P.A., Asher, D.M., Epstein, J.S., Kumar, S., 2009. Transfusion-transmitted babesiosis in the United States: summary of a workshop. Transfusion 49, 2759–2771.

Krause, P.J., 2019. Human babesiosis. International journal for parasitology 49, 165–174.

Leiby, D.A., 2011. Transfusion-transmitted *Babesia* spp.: bull’s-eye on *Babesia microti*. Clinical microbiology reviews 24, 14–28.

Liu, T., Zhang, L., Joo, D., Sun, S.C., 2017. NF-kappaB signaling in inflammation. Signal Transduct Target Ther 2, 17023-.

Lobo, C.A., Cursino-Santos, J.R., Alhassan, A., Rodrigues, M., 2013. *Babesia*: an emerging infectious threat in transfusion medicine. PLoS pathogens 9, e1003387.

Magni, R., Luchini, A., Liotta, L., Molestina, R.E., 2020. Proteomic analysis reveals pathogen-derived biomarkers of acute babesiosis in erythrocytes, plasma, and urine of infected hamsters. Parasitol Res 119, 2227–2235.

Mantel, P.Y., Hjelmqvist, D., Walch, M., Kharoubi-Hess, S., Nilsson, S., Ravel, D., Ribeiro, M., Gruring, C., Ma, S., Padmanabhan, P., Trachtenberg, A., Ankarklev, J., Brancucci, N.M., Huttenhower, C., Duraisingh, M.T., Ghiran, I., Kuo, W.P., Filgueira, L., Martinelli, R., Marti, M., 2016. Infected erythrocyte-derived extracellular vesicles alter vascular function via regulatory Ago2-miRNA complexes in malaria. Nat Commun 7, 12727.

Mantel, P.Y., Hoang, A.N., Goldowitz, I., Potashnikova, D., Hamza, B., Vorobjev, I., Ghiran, I., Toner, M., Irimia, D., Ivanov, A.R., Barteneva, N., Marti, M., 2013. Malaria-infected erythrocyte-derived microvesicles mediate cellular communication within the parasite population and with the host immune system. Cell Host Microbe 13, 521–534.

Mbagwu, S.I., Lannes, N., Walch, M., Filgueira, L., Mantel, P.Y., 2019. Human Microglia Respond to Malaria-Induced Extracellular Vesicles. Pathogens 9.

Nantakomol, D., Dondorp, A.M., Krudsood, S., Udomsangpetch, R., Pattanapanyasat, K., Combes, V., Grau, G.E., White, N.J., Viriyavejakul, P., Day, N.P., Chotivanich, K., 2011. Circulating red cell-derived microparticles in human malaria. The Journal of infectious diseases 203, 700–706.

Nogueira, P.M., Ribeiro, K., Silveira, A.C., Campos, J.H., Martins-Filho, O.A., Bela, S.R., Campos, M.A., Pessoa, N.L., Colli, W., Alves, M.J., Soares, R.P., Torrecilhas, A.C., 2015. Vesicles from different Trypanosoma cruzi strains trigger differential innate and chronic immune responses. J Extracell Vesicles 4, 28734.

Oz, H.S., Hughes, W.T., 1996. Acute fulminating babesiosis in hamsters infected with *Babesia microti*. International journal for parasitology 26, 667–670.

Piesman, J., Karakashian, S.J., Lewengrub, S., Rudzinska, M.A., Spielmank, A., 1986. Development of *Babesia microti* sporozoites in adult *Ixodes dammini*. International journal for parasitology 16, 381–385.

Regev-Rudzki, N., Wilson, D.W., Carvalho, T.G., Sisquella, X., Coleman, B.M., Rug, M., Bursac, D., Angrisano, F., Gee, M., Hill, A.F., Baum, J., Cowman, A.F., 2013. Cell-cell communication between malaria-infected red blood cells via exosome-like vesicles. Cell 153, 1120–1133.

Rudzinska, M.A., 1976. Ultrastructure of intraerythrocytic *Babesia microti* with emphasis on the feeding mechanism. J Protozool 23, 224–233.

Rudzinska, M.A., Trager, W., Lewengrub, S.J., Gubert, E., 1976. An electron microscopic study of *Babesia microti* invading erythrocytes. Cell Tissue Res 169, 323–334.

Silva, J.C., Cornillot, E., McCracken, C., Usmani-Brown, S., Dwivedi, A., Ifeonu, O.O., Crabtree, J., Gotia, H.T., Virji, A.Z., Reynes, C., Colinge, J., Kumar, V., Lawres, L., Pazzi, J.E., Pablo, J.V., Hung, C., Brancato, J., Kumari, P., Orvis, J., Tretina, K., Chibucos, M., Ott, S., Sadzewicz, L., Sengamalay, N., Shetty, A.C., Su, Q., Tallon, L., Fraser, C.M., Frutos, R., Molina, D.M., Krause, P.J., Ben Mamoun, C., 2016. Genome-wide diversity and gene expression profiling of *Babesia microti* isolates identify polymorphic genes that mediate host-pathogen interactions. Scientific reports 6, 35284.

Silverman, J.M., Clos, J., de’Oliveira, C.C., Shirvani, O., Fang, Y., Wang, C., Foster, L.J., Reiner, N.E., 2010a. An exosome-based secretion pathway is responsible for protein export from Leishmania and communication with macrophages. Journal of cell science 123, 842–852.

Silverman, J.M., Clos, J., Horakova, E., Wang, A.Y., Wiesgigl, M., Kelly, I., Lynn, M.A., McMaster, W.R., Foster, L.J., Levings, M.K., Reiner, N.E., 2010b. Leishmania exosomes modulate innate and adaptive immune responses through effects on monocytes and dendritic cells. J Immunol 185, 5011–5022.

Skariah, S., Arnaboldi, P., Dattwyler, R.J., Sultan, A.A., Gaylets, C., Walwyn, O., Mulhall, H., Wu, X., Dargham, S.R., Mordue, D.G., 2017. Elimination of *Babesia microti* Is Dependent on Intraerythrocytic Killing and CD4(+) T Cells. J Immunol 199, 633–642.

Terkawi, M.A., Cao, S., Herbas, M.S., Nishimura, M., Li, Y., Moumouni, P.F., Pyarokhil, A.H., Kondoh, D., Kitamura, N., Nishikawa, Y., Kato, K., Yokoyama, N., Zhou, J., Suzuki, H., Igarashi, I., Xuan, X., 2015. Macrophages are the determinant of resistance to and outcome of nonlethal *Babesia microti* infection in mice. Infection and immunity 83, 8–16.

Thekkiniath, J., Kilian, N., Lawres, L., Gewirtz, M.A., Graham, M.M., Liu, X., Ledizet, M., Ben Mamoun, C., 2019. Evidence for vesicle-mediated antigen export by the human pathogen *Babesia microti*. Life Sci Alliance 2.

Vannier, E.G., Diuk-Wasser, M.A., Ben Mamoun, C., Krause, P.J., 2015. Babesiosis. Infectious disease clinics of North America 29, 357–370.

Yanez-Mo, M., Siljander, P.R., Andreu, Z., Zavec, A.B., Borras, F.E., Buzas, E.I., Buzas, K., Casal, E., Cappello, F., Carvalho, J., Colas, E., Cordeiro-da Silva, A., Fais, S., Falcon-Perez, J.M., Ghobrial, I.M., Giebel, B., Gimona, M., Graner, M., Gursel, I., Gursel, M., Heegaard, N.H., Hendrix, A., Kierulf, P., Kokubun, K., Kosanovic, M., Kralj-Iglic, V., Kramer-Albers, E.M., Laitinen, S., Lasser, C., Lener, T., Ligeti, E., Line, A., Lipps, G., Llorente, A., Lotvall, J., Mancek-Keber, M., Marcilla, A., Mittelbrunn, M., Nazarenko, I., Nolte-’t Hoen, E.N., Nyman, T.A., O’Driscoll, L., Olivan, M., Oliveira, C., Pallinger, E., Del Portillo, H.A., Reventos, J., Rigau, M., Rohde, E., Sammar, M., Sanchez-Madrid, F., Santarem, N., Schallmoser, K., Ostenfeld, M.S., Stoorvogel, W., Stukelj, R., Van der Grein, S.G., Vasconcelos, M.H., Wauben, M.H., De Wever, O., 2015. Biological properties of extracellular vesicles and their physiological functions. J Extracell Vesicles 4, 27066.

